# Corticospinal excitability is influenced by the recent history of electrical digital stimulation: implications for the relative magnitude of short-latency afferent inhibition

**DOI:** 10.1101/2022.02.02.478861

**Authors:** Marie Trolle Bonnesen, Søren Asp Fuglsang, Hartwig Roman Siebner, Lasse Christiansen

## Abstract

**Background:** Electrical stimulation of the hand can suppress the motor evoked potential (MEP) elicited by transcranial magnetic stimulation (TMS) of the primary motor hand area (M1-HAND) when the afferent stimulus arrives in M1-HAND at the time of TMS. The magnitude of short-latency afferent inhibition (SAI) is expressed as the ratio between the conditioned and unconditioned MEP and is widely used to probe sensorimotor interactions in human physiological studies.

**Objective/Hypothesis:** We hypothesized that corticospinal excitability and SAI are influenced by the recent history of peripheral electrical stimulation.

**Methods:** In twenty healthy participants, we recorded MEPs from the right first dorsal interosseus muscle. MEPs were evoked by single-pulse TMS of the left M1-HAND alone (unconditioned TMS) or by TMS preceded by electrical stimulation of the right index finger (“homotopic” conditioning) or little finger (“heterotopic” conditioning). The three conditions were pseudo-randomly intermixed or delivered in blocks in which a single condition was repeated five or ten times. MEP amplitudes and SAI magnitudes were compared using linear mixed effect models.

**Results:** All stimulation protocols consistently produced SAI, which was stronger after homotopic stimulation. Randomly intermingling the three stimulation conditions reduced the relative magnitude of homotopic and heterotopic SAI as opposed to blocked stimulation. The apparent attenuation of SAI was caused by a suppression of the unconditioned but not the conditioned MEP amplitude during the randomly intermixed condition.

**Conclusion(s):** The recent history of afferent stimulation modulates corticospinal excitability. This “history effect” impacts on the relative magnitude of SAI depending on how conditioned and unconditioned responses are intermixed and needs to be taken into consideration when probing afferent inhibition and corticospinal excitability.

## Introduction

Humans display extraordinary dexterous control of the hand and digits. In the face of the rapidly changing sensory environment, fine motor control of the fingers depends on the ability of the central nervous system to rapidly integrate somatosensory inputs from the upper limb into precise motor outputs [1,2]. Efficient integration needs to take into account both past and present sensory inputs [3]. The extent to which dynamic changes in afferent inputs from the hand shape motor output is still largely unexplored.

Fast cortical integration of somatosensory inputs from the hand can be studied by measuring short-latency afferent inhibition (SAI) using a conditioning-test paradigm [4]. A conditioning peripheral electrical stimulus produces a highly synchronized afferent sensory volley from the hand. This afferent input markedly inhibits the motor evoked potentials (MEP) evoked by transcranial magnetic stimulation (TMS) of the contralateral motor hand area (M1-HAND), if the TMS pulse is given a few milliseconds after the arrival of the afferent volley in the pericentral sensorimotor cortex [4]. SAI is suggested to be mediated either by a direct thalamocortical pathway or through a short relay in the primary sensory cortex [5,6]. SAI arises in M1-HAND, because the SAI conditioning-test paradigm results in a suppression of late TMS-evoked I-waves [4]. The magnitude of SAI displays somatotopy. “Homotopic” afferent stimulation that secures spatial correspondence between the peripherally stimulated afferents and the trans-cranially stimulated hand muscle results in stronger SAI than “heterotopic” afferent stimulation with a somatotopic mismatch between the stimulated afferents and hand muscle [7–9]. Over the past two decades SAI has been used as a tool to explore and advance the understanding of sensorimotor integration at the level of the sensorimotor cortex. SAI has also been widely used to investigate altered signaling and probe changes in sensorimotor function in neurodegenerative diseases such as Parkinson’s disease [10].

SAI is commonly expressed as the ratio between MEP amplitudes evoked by conditioned and unconditioned TMS pulses, allowing for comparisons between different states and groups with unequal corticomotor excitability. Reports of SAI measured from the first dorsal interosseus (FDI) in healthy humans range from 20-60% inhibition despite obtained with comparable parameter settings [11]. The varying magnitude of SAI may reflect intrinsic fluctuations in sensorimotor excitability and may be influenced by how conditioned and unconditioned responses are intermixed. This notion is corroborated by previous studies demonstrating that single-pulse MEP amplitudes are modulated by interstimulus intervals suggestive of long lasting effects of previous stimulation in the order of seconds [12,13]. Furthermore, previous findings show that SAI interacts with other excitatory and inhibitory intracortical circuitries [14,15], suggesting that the inhibitory effect of afferent inputs to M1-HAND may depend on the “excitability state” of other cortical but possibly also spinal circuits that control corticomotor output. In this study, we tested the hypothesis that the relative magnitude of SAI is influenced by the way in which afferent conditioned and unconditioned stimulation are intermixed. To this end, we manipulated the sensorimotor context by delivering homo- and heterotopically conditioned and unconditioned TMS stimuli either randomly intermixed or repeatedly in blocks of five or ten identical stimuli.

## Material and methods

### Participants

Twenty healthy volunteers (mean age 24.9, ± 3.7 SD, nine women) participated in the study. All participants were right handed, assessed by the Edinburgh Handedness Inventory [16], and had no history of neurological disease. Participants were screened for contraindications to MRI and TMS [17] and gave written, informed consent to the experiment. The experiment received ethical approval from the Ethics Committee of Capital Region of Denmark (No. H-17023857) and complied with the Declaration of Helsinki.

### Experimental design

The experiment was designed to investigate if the relative magnitude of SAI in FDI differs, depending on how peripherally conditioned TMS pulses are intermingled with unconditioned TMS pulses.

On the day of the main experiment, participants were seated comfortably in a chair, looking straight ahead with their left arm on a pillow and right arm on a table with a foam cushion. The head was supported by a headrest and participants were requested to relax with the eyes open. We recorded series of sixty MEPs from the FDI muscle with an intertrial interval (ITI) of 5.5 sec with 20% jitter. Each series consisted of three experimental stimulation conditions (Fig. 1A): 1) TMS delivered to left M1-HAND alone (unconditioned, *TS*), 2) TMS preceded by electrical stimulation of the right index finger (conditioned, *D2+TS*), or 3) TMS preceded by electrical stimulation of the right little finger (conditioned, *D5+TS*). Each series of sixty MEPs was delivered in one of three stimulation patterns (Fig. 1B): 1) stimulation conditions were *pseudo*-randomly intermixed but limited to five successive stimulations within the same condition (*Random*), 2) stimulation conditions were delivered in blocks in which a single condition was repeated five times (*B5*), or 3) stimulation conditions were delivered in blocks in which a single condition was repeated ten times (*B10*). In total, we recorded six series of sixty MEPs so that each pattern was recorded twice. By doing so, we recorded a total of forty MEPs for each condition in each pattern. The order of patterns and conditions in *B5* and *B10* was balanced across participants with a fixed order within a participant.

**Fig. 1.**
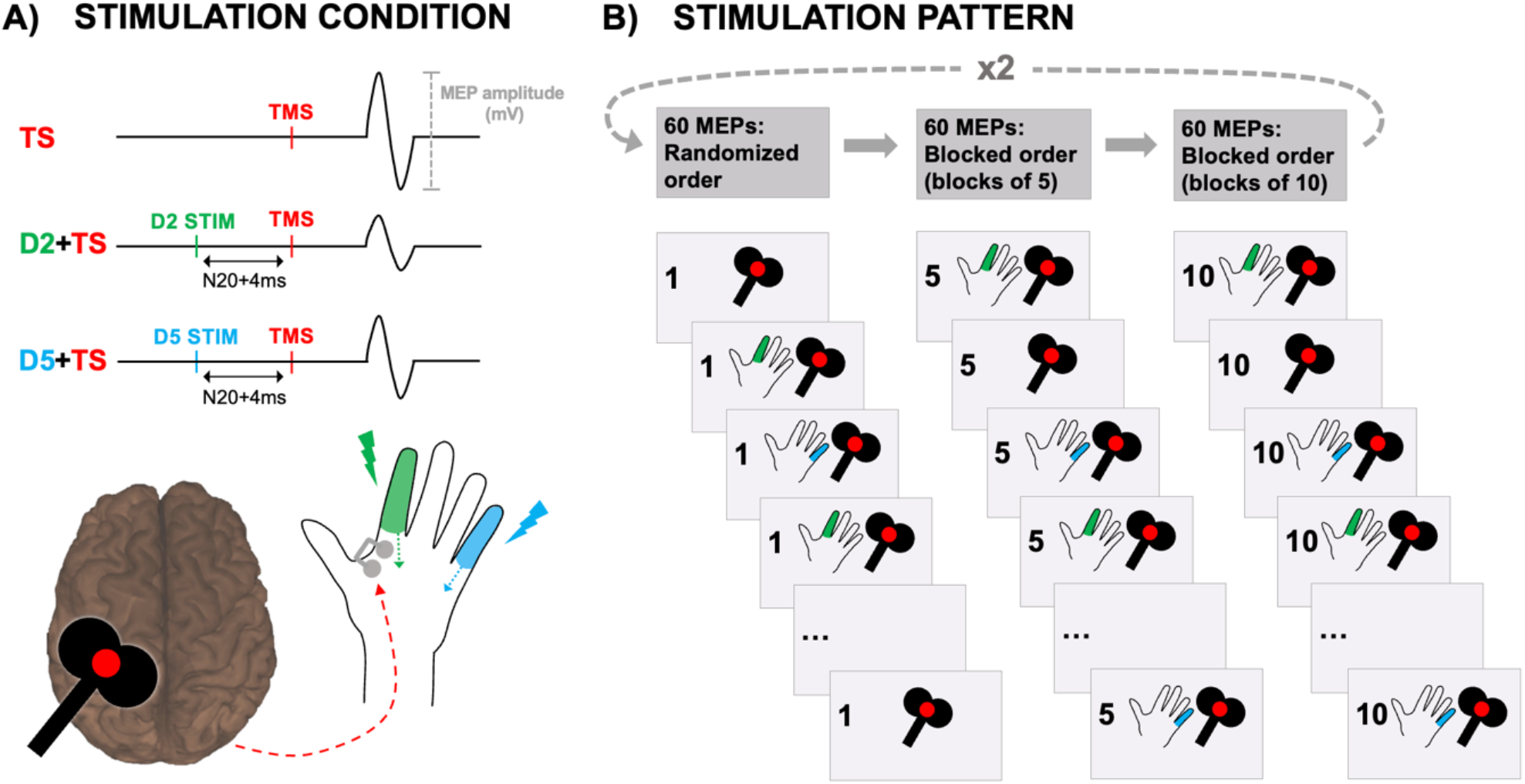
Experimental design. **A)** Motor evoked potentials (MEPs) were recorded from surface electrodes (grey dots). Participants were stimulated with three different stimulation conditions combining TMS at the FDI hotspot (red dot) with peripheral electrical stimulation of the right index (D2, green) or little finger (D5, blue): 1) Unconditioned TMS alone (*TS*), 2) Conditioned TMS and D2 stimulation (*D2+TS*), 3) Conditioned TMS and D5 stimulation (*D5+TS*). **B)** The three stimulation conditions were combined in three different stimulation patterns: 1) 60 MEPs with randomized order of stimulation conditions, 2) 60 MEPs with stimulation conditions repeated in blocks of five MEPs, 3) 60 MEPs with stimulation conditions repeated in blocks of ten MEPs. The stimulation pattern on the figure is an example for one subject.

### Neuronavigated transcranial magnetic stimulation

Participants were co-registered to their individual T1-weighted MRI scan (Magnetom Prisma 3T, Siemens), acquired prior to the day of testing, using a frameless neuronavigation system (*Localite*, GmbH, Bonn, Germany). A trained investigator manually placed a stimulation target on the lip of the precentral gyrus in the left M1-HAND [18].

Single, biphasic pulses inducing AP-PA currents orthogonal to the central sulcus were delivered through a Cool-B35 HO butterfly figure-of-eight coil connected to a MagPro X100 with MagOption stimulator (Magventure, Skovlunde, Denmark). Small corrections of the coil position and orientation were made to find the physiological hotspot of the FDI muscle yielding the largest MEP amplitude. Precise positioning of the TMS coil relative to the participant’s head was continuously monitored. Finally, the TMS stimulation intensity was adjusted to reach a mean *TS* MEP amplitude of 0.5-1.0 mV.

### Conditioning peripheral electrical stimuli

Peripheral electrical stimuli were given to the right index and little finger through bipolar ring electrodes positioned with the cathode proximal and anode distal to the proximal interphalangeal joint. Single square pulses of 200 μs duration (Digitimer stimulator, model DS7A and DS7AH) were applied with an intensity of three times perceptual threshold [4]. Participants did not experience the stimulus as painful.

In order to elicit SAI, the interstimulus interval (ISI) between the TMS pulse and the peripheral electrical stimulation was determined relative to the individual latency of the N20 component of the somatosensory evoked potential (SSEP) evoked by electrical stimulation of the index finger plus 4ms [19]. To record SSEPs on the scalp, the reference electrode was places at the Fz, and the active electrode was placed at the CP3, 10% posterior to the C3 of the 10-20 EEG system [20]. The N20 peak was identified from the average trace of two to six hundred responses.

### Electromyographic recording

The electromyography (EMG) activity was recorded from the relaxed right FDI with Ag/AgCl surface electrodes (Ambu Neuroline 700, Columbia, USA) in a belly-tendon montage. The active electrode was placed over the motor point of the right FDI muscle, the reference electrode on the metacarpophalangeal joint of the right index finger and the ground electrode on the head of the right ulnar bone. The signal was amplified (1000x, Digitimer Ltd, D360), bandpass-filtered (5Hz-2KHz), digitized and stored on a computer for offline analysis (Signal Version 4.11, Cambridge Electronic Design, Cambridge, UK). EMG activity 100ms prior to peripheral electrical stimulation was monitored online, and participants were instructed and trained in maintaining full relaxation of the FDI muscle. All EMG recordings were visually inspected and trials with voluntary background contraction were removed. For each trial the peak-to-peak MEP amplitude in the time window between 20ms and 40ms after the TMS pulse was extracted. In the trial-by-trial exploration of MEP amplitudes within the *Random* stimulation pattern, the first trial was always excluded as it was not influenced by PREVIOUS CONDITION resulting in a total of 112 to 118 trials per subjects.

### Data analysis

Mean peak-to-peak MEP amplitudes for each stimulation condition in each pattern were computed for participants separately. In addition, the relative magnitude of SAI was calculated as the ratio between the conditioned and unconditioned responses. Lastly, for trial-by-trial analyses, MEP amplitudes of single trials were used to model the effect of the previous condition on subsequent responses within the randomized pattern.

Statistical inferences were based on linear mixed-effects models (LMMs) and performed in R Studio Version 1.4.1717 (RStudio Team, 2021), running on R Version 4.1.1 (R Core Team, 2021), with the ggplot2 [21], lme4 [22] and lmerTest [23] packages. When incorporating single trial MEP amplitudes, the amplitudes were log-transformed before modelling to reduce skewness in the data. All data are visually displayed in the original form (non-transformed) for clarity. F- and p-values for the mixed models were computed with Type-III analysis of variance using Satterthwaite’s method to estimate degrees of freedom [23]. The significance threshold for null hypothesis testing was set at p<0.05. The p-values of post hoc pairwise comparisons were adjusted for multiple comparisons using the Bonferroni correction.

Effects of CONDITION and PATTERN on mean MEP amplitude were explored with LMMs including intercepts for ‘subjects’ as a random factor. A similar analysis was conducted to investigate relative suppression for conditioned responses.

Effects of PREVIOUS CONDITION for data acquired in the *Random* pattern were interrogated using LMMs fit to data from individual trials for all subjects. Separate models were fit to data from *TS*, *D2+TS* and *D5+TS* conditions. Each model included PREVIOUS CONDITION as fixed effects and ‘subject’ as a random factor. To better distinguish effects of previous stimulation condition from the amplitude of the previous MEP, the models further included separate regressors of no interest for each subject and each condition (centered within condition and subject) reflecting specific effects of PREVIOUS AMPLITUDE.

Trial-by-trial effects of PREVIOUS AMPLITUDE were further probed by estimating the Pearson’s correlation between response amplitudes and the preceding response amplitude (i.e., lagged correlations) for data acquired in the *Random* pattern. This was done separately for each subject within each transition (e.g., *TS* to *TS*). The coefficients were transformed using Fisher’s Z-transformation, averaged across subjects and inverse transformed using the inverse Fisher’s Z-transformation to yield a group-mean estimate. Furthermore, this same process was repeated over 50000 iterations, but in each iteration with phase scrambled response vectors to obtain a phase scrambled noise distribution. Randomization testing was applied to reveal differences between the computed and phase scrambled lagged correlations.

## Results

Single-pulse TMS was applied to M1-HAND with a mean stimulation intensity of 71.4% maximal stimulator output (SD: 11.6%). Mean peripheral stimulus intensity was 13.95 mA (SD: 3.27 mA) for the right index finger and 13.53 mA (SD: 3.21 mA) for the right little finger. Mean N20 latency of cortical SSEP was 22.2 ms (SD: 1.2ms). None of the participants reported any adverse effects during the experiment.

### Effect of stimulation condition and pattern on mean MEP amplitudes

All subjects displayed SAI regardless of the stimulation pattern. Inhibition was stronger following homotopic as opposed to heterotopic afferent stimulation (Fig. 2A). In agreement with our hypothesis, the temporal pattern, at which stimulation conditions were applied, had an influence on corticospinal excitability during SAI measurements (Fig.2). MEP amplitudes evoked by the unconditioned test stimulus alone were consistently attenuated, when data acquisition employed a *Random* pattern as opposed to a blocked (*B5* or *B10*) stimulation pattern (Fig.2B). In contrast, MEP amplitudes evoked by peripherally conditioned test stimuli showed a comparable mean MEP amplitude during *Random* and *Blocked* stimulation (Fig.2C,2D).

**Fig. 2.**
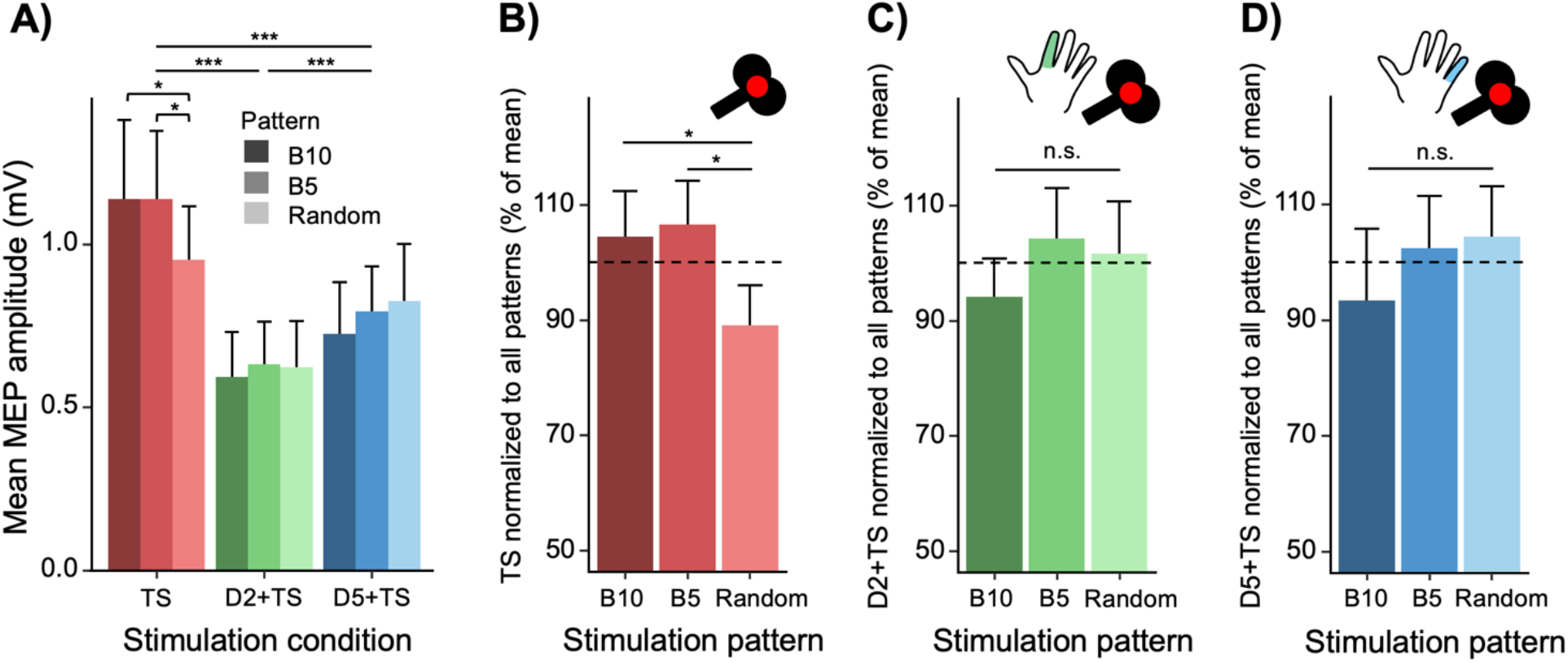
Group data of the MEP amplitudes in all stimulation conditions and patterns. **A)** Group mean MEP amplitudes of all three stimulation conditions (*TS, D2+TS, D5+TS*) divided into each stimulation pattern (*Random, B5* and *B10*). Significant differences (***) in the MEP amplitude between all the three stimulation conditions across patterns. Furthermore, significant smaller *TS* amplitude in *Random* compared to *B5* (*) and *B10* (*). **B-D)** Explication of CONDITION by PATTERN interaction for illustrative purposes. MEP amplitudes for each stimulation condition normalized to the grand mean across patterns within each condition. Bars representing group mean + 95% CI. *** = p<0.001, * = p<0.05. MEP = motor evoked potential, TS = test stimulus (TMS alone), D2+TS = TMS preceded by index finger stimulation, D5+TS = TMS preceded by little finger stimulation, Random = randomized pattern, B5 = stimulation in blocks of five MEPs, B10 = stimulation in blocks of ten MEPs.

These findings were statistically confirmed by the LMM revealing a main effect of CONDITION (F_(2,152)_=81.42, p<0.001), a CONDITION x PATTERN interaction effect (F_(4,152)_=3.058, p=0.019), but no main effect of PATTERN (F_(2,152)_=1.172, p=0.313). Post-hoc comparisons within CONDITION showed significantly smaller responses following *D2+TS* as compared to *D5+TS* and *TS*, as well as for *D5+TS* compared to *TS* (all p<0.001) (Fig. 2A). Furthermore, significantly smaller MEP amplitudes were evoked by the *TS* condition during the *Random* pattern compared to the blocked-*B5* (p=0.011) and blocked-*B10* (p=0.011) patterns (Fig. 2A,B). The *Random* pattern did not induce a relative suppression of MEP amplitude of the conditioned responses compared to a blocked stimulation pattern (p>0.35, Fig. 2A,C,D).

Taken together, results revealed a relative suppression of unconditioned *TS* responses during a *Random* pattern compared to *Blocked* patterns of stimulation. In contrast, MEP amplitudes in the context of SAI evoked by *D2+TS* or *D5+TS* stimulation were unaffected by the pattern of stimulation.

### Relative magnitude of SAI

The suppression of the unconditioned MEP amplitude during the *Random* stimulation pattern resulted in a relative attenuation of SAI compared to the two blocked-*B5* and *B10* pattern (Fig. 3). When using the relative MEP size (*D2+TS* or *D5+TS* expressed as percentage of the unconditioned *TS* response) as depended variable, the LMM showed a significant main effect of PATTERN (F_(2,95)_=12.88, p<0.001) but no significant CONDITION x PATTERN interaction (F_(2,95)_=0.448, p=0.640). Pairwise comparisons of PATTERN revealed a relative attenuation of SAI (i.e. less inhibition indexed as percentage of unconditioned *TS* response) when stimulation conditions were randomized compared to blocked conditions (both p<0.001), displaying that the relative magnitude of afferent inhibition was reduced during the *Random* pattern. The LMM also confirmed that the relative magnitude of SAI was stronger for homotopic stimulation (*D2+TS*) compared to heterotopic stimulation (*D5+TS*) evident as a main effect of CONDITION (F_(1,95)_=29.99, p<0.001).

**Fig. 3.**
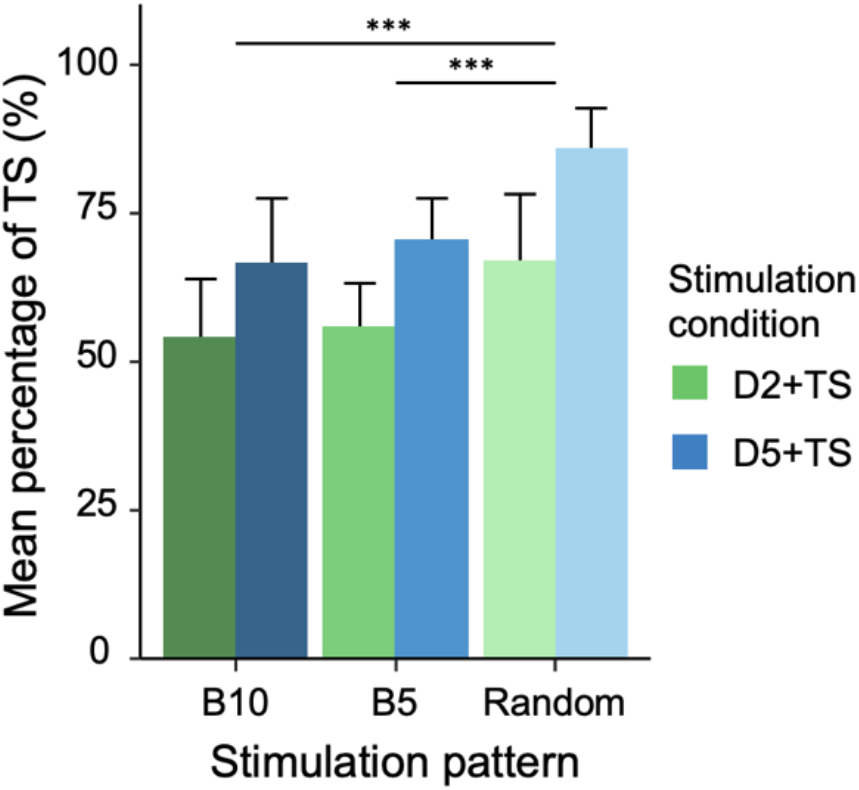
Relative magnitude of SAI. Relative magnitude of SAI in the different stimulation conditions and patterns. Mean MEP amplitude of the two conditioned responses (*D2+TS* and *D5+TS*) expressed as percentage of mean MEP amplitude of *TS*. Less SAI in stimulation pattern *Random* compared to *B5* (***) and *B10* (***). Furthermore, less SAI in stimulation condition *D5+TS* compared to *D2+TS* (***). Bars representing group mean + 95% CI. *** = p<0.001. MEP = motor evoked potential, TS = test stimulus (TMS alone), D2+TS = TMS preceded by index finger stimulation, D5+TS = TMS preceded by little finger stimulation, Random = pseudo randomized pattern, B5 = stimulation in blocks of five MEPs, B10 = stimulation in blocks of ten MEPs.

### “Recency effects” in the randomized stimulation pattern

In a follow-up analysis, we explored the possibility that lower average *TS* MEP amplitude found during a *Random* pattern of stimulation was caused by a suppression from the preceding trial. We tested for two types of “recency effects” from the antecedent trial on the MEP amplitude in the following trial, namely effects related to the previous stimulation condition and effects related to the MEP size elicited in the previous trial.

Only the unconditioned *TS* showed a recency effect of the previous stimulation condition on MEP amplitude in the subsequent trial. MEP amplitudes evoked by the unconditioned *TS* were smaller when preceded by a heterotopic stimulation trial (Fig. 4A). Trial-by-trial analysis of log-transformed *TS*-only amplitudes employed a conservative LMM modelling PREVIOUS CONDITION with the effects of PREVIOUS AMPLITUDE regressed out on a subject and condition specific level. The LMM yielded a significant main effect of PREVIOUS CONDITION (F_(2,721.92)_=7.68, p<0.001) (Fig. 4A). Post-hoc comparisons of PREVIOUS CONDITION showed that unconditioned MEP amplitudes were reduced following a heterotopic *D5+TS* conditioned trial compared to an unconditioned *TS* trial (p<0.001). However, no difference was found when the preceding trial was a homotopic *D2+TS* compared to an unconditioned *TS-only* trial (p=0.246), or when the preceding trial was a heterotopic compared to homotopic conditioning stimulation (p=0.070). As expected from the lack of differences between conditioned trials obtained with blocked versus randomized pattern, no recency effects were evident in conditioned MEPs (Fig. 4B and 4C). The two LMMs showed no main effect of PREVIOUS CONDITION (present condition *D2+TS*: F_(2,731.71)_=1.912, p=0.148; present condition *D5+TS*: F_(2,711.17)_=1.009, p=0.365).

**Fig. 4.**
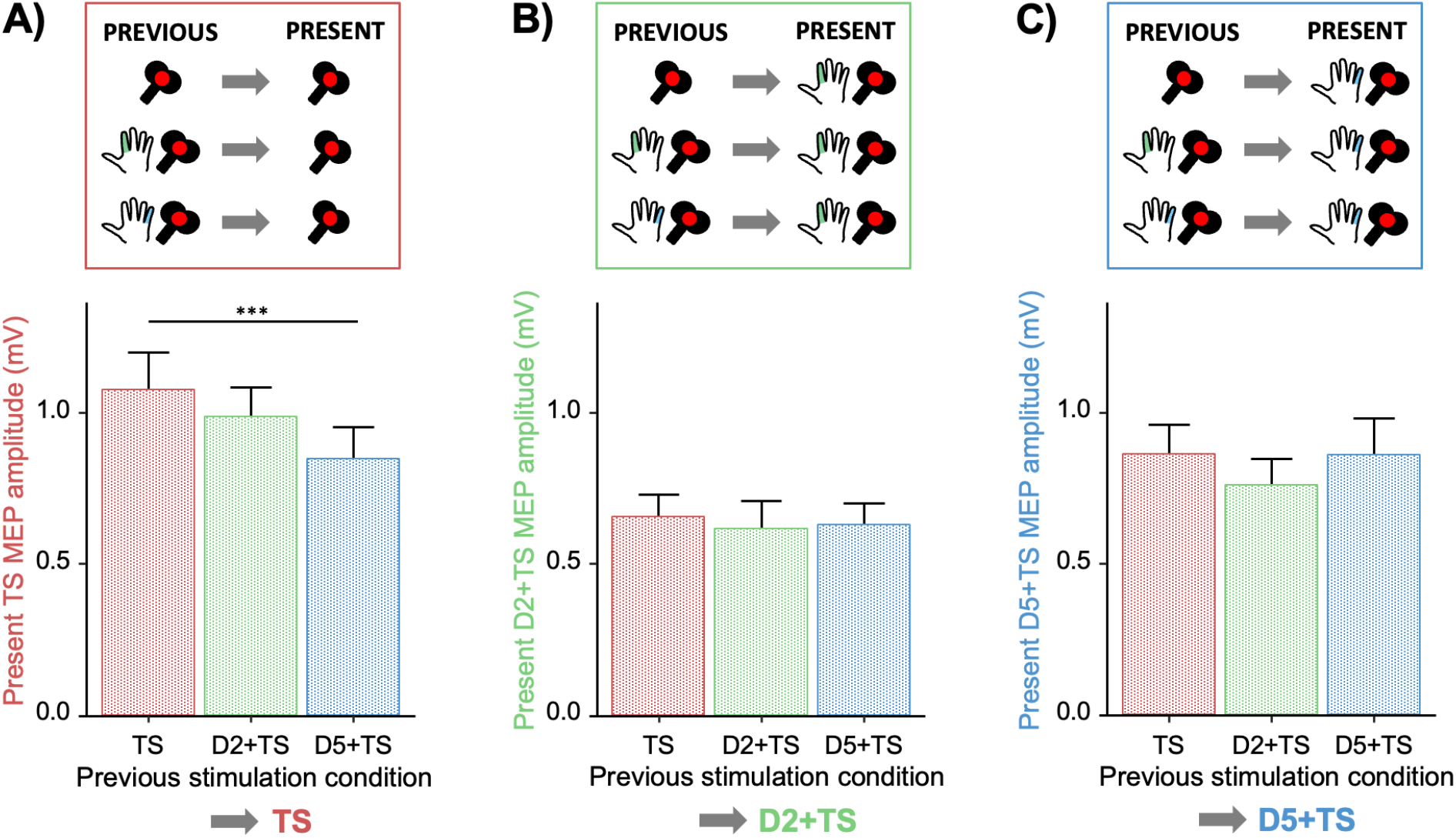
“Recency effects” in the randomized pattern. **A)** Effect of recent (previous) stimulation condition on the present MEP amplitude of *TS*. Significant effect of PREVIOUS CONDITION (***) with smaller *TS* MEP amplitude when the previous condition was *D5+TS* compared to *TS* (***). **B)** Effect of PREVIOUS CONDITION on the present MEP amplitude of *D2+TS*. No effect of previous condition. **C)** Effect of PREVIOUS CONDITION on the present MEP amplitude of *D5+TS*. No effect of previous condition. Bars representing group mean + 95% CI. *** = p<0.001. MEP = motor evoked potential, TS = test stimulus (TMS alone), D2+TS = TMS preceded by index finger stimulation, D5+TS = TMS preceded by little finger stimulation.

Associations between MEP amplitudes elicited in each trial and MEP amplitudes elicited in the preceding trial were further explored across all three conditions within the *Random* stimulation pattern, revealing a positive relationship extending back all three lags investigated (see supplementary figure 1, mean rho for lag 1=0.20, lag 2=0.12 and lag 3=0.09, p<0.01, two-sided randomization tests, Bonferroni corrected for three comparisons). Due to the exponential increase in number of combinations of present and antecedent trials when extended the analysis more than one lag, we restricted further analyses to the first lag interrogating the effect on present *TS* amplitudes.

To further dissociate the positive correlation for the first lag (i.e. between MEP amplitude and preceding MEP amplitude) from the effect of previous stimulation condition, correlations were explored separately for all three previous conditions with *TS* as the present condition (e.g., for *TS* trials preceded by *D2+TS* trials, see supplementary figures S2-S4). MEP amplitude correlated positively with the amplitude of the previous amplitude in *TS* trials preceded by *TS* (p=0.005) and *D2+TS* trials (p<0.001), but not in *TS* trials preceded by *D5+TS* trials (p=0.52, two-sided randomization tests, Bonferroni corrected for nine comparisons). The effect of previous amplitude partly generalized to the two conditioned stimulation conditions (corrected p-values range <0.001-0.586, see supplementary figures S5-S10).

For internal validation, similar analyses were conducted for *TS-TS* combination within the blocked-*B5* and blocked-*B10* patterns excluding the first ‘transition’ trial. The analyses confirmed the results from the *Random* pattern in that the amplitude of the preceding MEP affected the amplitude of the present MEP within the *TS* blocks of 5 (p=0.001) and 10 (p=0.018), Bonferroni corrected for two comparisons).

Together, the results of the follow-up analyses showed that when accounting for the effect of previous MEP amplitude, the randomized stimulation pattern displayed a recency effect where the previous stimulation condition impacted the consecutive unconditioned but not conditioned trials. Furthermore, a generalized positive effect of previous MEP amplitude was visible albeit more pronounced in *TS* and *D2+TS* conditions.

## Discussion

We found that peripherally conditioned TMS pulses intermingled with unconditioned TMS pulses led to a reduction in corticospinal excitability expressed as attenuated motor responses to unconditioned TMS pulses. The attenuated MEP response to single-pulse stimulation reduced the relative magnitude of SAI when conditions were pseudo-randomly intermixed. In contrast, the absolute magnitude of SAI, reflected by the peripherally conditioned MEP amplitudes, remained stable regardless of stimulation pattern. Furthermore, we demonstrated that the recent “history of sensorimotor stimulation” in the previous trial tuned the corticospinal excitability in the next trial. We also replicated a spatial gradient of SAI with stronger SAI after homotopic relative to heterotopic electrical finger stimulation.

### SAI depends on the stimulation pattern

Our main finding was that the relative magnitude of SAI decreases when stimulation conditions are intermixed pseudo-randomly as opposed to a blocked pattern of stimulation consisting of five or ten of the same condition. Critically, the peripherally conditioned MEP amplitude was comparable during random and blocked stimulation, showing that SAI was comparable among stimulation patterns. The apparent reduction in the relative SAI magnitude was entirely driven by a reduction in MEP amplitude of the unconditioned MEP response. Together, the results indicate that the sensorimotor context, experimentally introduced by the temporal pattern of stimulation conditions, influenced corticospinal excitability, causing a relative suppression of corticospinal excitability when intermingling afferent conditioned and unconditioned trials. This context-dependent attenuation in corticospinal excitability caused an apparent attenuation of the relative magnitude of SAI. Our findings are highly relevant from a methodological perspective, as most studies express the magnitude of SAI in relation to the unconditioned MEP response. We infer that future studies should consider both the relative and absolute strength of SAI, factoring in the corticospinal excitability level during SAI measurements. The results also call for future research into the dynamic regulation of corticospinal excitability based on the recent history of sensorimotor stimulation. The dynamic sensorimotor interplay may be altered in neurodegenerative diseases such as Alzheimer’s disease or Parkinson’s disease, that display alterations in SAI [10]. This may yield novel insights into the neural pathways as well as the involved neurotransmitters of the recency effect.

While, to the best of our knowledge, the effect of the sensorimotor stimulation pattern on the relative magnitude of SAI has not been shown before, previously reported magnitudes of SAI vary greatly across studies [11]. We speculate, that this variation across studies may in part be caused by different ways of combining conditioned and unconditioned stimuli during the assessment of SAI. We would like to point out that the stimulation pattern is not the only factor that contributes to the magnitude of inhibition. Other contributing factors include stimulation settings such as peripheral stimulation intensity [24], the ISI [8] and the intensity of the TMS pulse [25,26], as well as individual biological factors such as age [27,28], handedness [29] and genotype [30].

### Corticospinal excitability changes with sensorimotor context

The attenuation of the relative magnitude of SAI was solely caused by a relative suppression of the unconditioned MEP response, when the peripherally conditioned and unconditioned TMS trials were intermixed. Rapid context-dependent modulations of the MEP amplitude are a well-known feature in MEP studies. A slight tonic contraction of the target muscle leads to a marked facilitation of the MEP relative to the MEP evoked during muscle relaxation [31,32]. The mean MEP amplitude can also display rapid modulations at rest. For instance, the power or phase of ongoing cortical oscillatory activity is associated with differences in corticospinal excitability, causing relative differences in mean MEP amplitudes [33,34]. The suppression of the unconditioned test response when intermixed with conditioned stimuli may therefore originate from fluctuations in sensori-cortico-motor excitability caused by dynamic changes in sensorimotor stimulation.

### The recent trial matters

Previous TMS studies, which have given unconditioned single TMS pulses at a slow repetition rate, reported systematic changes in MEP amplitude within and between blocks of stimulation [12,13], indicating that the repeated administration of TMS can produce relative shifts in corticospinal excitability even at relatively long interstimulus intervals. Our study extends these findings by showing that the MEP amplitude of the previous trial is positively correlated with the MEP amplitude of the next trial. We infer that even at an ISI of several seconds, the corticospinal excitability of the previous trial “spills over” to the next trial.

Our analyses revealed another type of recency effect of the preceding trial on the MEP response evoked in the consecutive trial in the pseudo-randomized stimulation pattern. We found that the sensorimotor context created by the previous trial impacted the unconditioned MEP amplitude in the next trial. When accounting for the general effect of the previous MEP amplitude on the present MEP response, the mean MEP amplitudes were reduced, if the previous trial included a peripheral nerve stimulation. Of note, this recency effect was only expressed when an unconditioned TMS pulse was given in the following trial, but the preceding sensorimotor stimulation had no effect when the next trial also involved an afferent conditioning stimulus. This selective recency effect of sensorimotor stimulation on corticospinal excitability, indicates that the recent combination of sensory input from the contralateral hand and central stimulation of M1 impacts the excitability of the corticomotor output several seconds later. We argue that the selective suppressive recency effect of sensorimotor stimulation accounted for the mean reduction in corticospinal excitability during randomized stimulation and led to an apparent attenuation of SAI. Our results therefore show that a conditioned stimulation produces not only a prominent inhibition at short latency (SAI), but also causes longer lasting inhibition of the corticospinal output which might decay in the order of seconds.

We can only speculate about the physiological mechanisms that underpin the recency effect related to recent afferent stimulation of the contralateral hand. The preceding sensorimotor stimulation may suppress corticospinal excitability at the spinal level or via thalamocortical, intracortical or cerebellar pathways. The interaction between SAI and intracortical excitatory [14] and inhibitory [15] interneural circuitries known to shape corticospinal output, supports the notion that inputs to M1 from upstream pre-motor areas, as well as thalamocortical and corticocortical inputs known to influence pyramidal output and corticomotor excitability, may interact with the motor output. SAI has been shown to inhibit other paired-pulse paradigms such as short interval intracortical inhibition (SICI) [15] and long-interval cortical inhibition (LICI) [26], and facilitate short interval intracortical facilitation (SICF) [14]. The above findings demonstrate that SAI not only affect layer 5 corticospinal neurons directly but also intracortical connections, suggesting that the suppression of corticospinal excitability in the trial following a conditioned response could be due to interactions at the cortical level. However, while SAI predominantly reflects activity in a transcortical reflex loop [35], afferent electrical stimulation of a digit also elicits reflex responses of spinal origin [36–38]. It is yet to be demonstrated that such reflex activity in the spinal circuitry can suppress the motoneuronal response to descending activation several seconds later, but a spinal contribution to the context-dependency cannot be excluded. Future studies are needed to clarify which networks that mediate the observed context-dependency and tune corticospinal excitability to the recent history of sensory inputs. Studying patients with cerebellar or thalamic lesions, probing changes in oscillatory activity in the order of seconds with EEG, or examining the effect on subcortically evoked test responses could contribute to elucidate the neural mechanisms underlying the inhibitory recency effect occurring when unconditioned responses are preceded by conditioned stimulation.

### The spatial gradient of SAI is reversed for sensorimotor recency effects

We found a spatial gradient of SAI regardless of the stimulation pattern, as the inhibitory effect of afferent stimulation was more pronounced with homotopic (index finger) electrical stimulation compared to heterotopic (little finger) electrical stimulation. The stronger expression of SAI with homotopic stimulation is in line with previous results, showing either a weaker afferent inhibition after heterotopic stimulation [7,8] or even a heterotopic facilitation [9]. As SAI has been shown to be mediated at the level of the cortex [4], the results show that a peripherally evoked afferent volley inhibits the excitability of the motor cortex in a somatotopic fashion, being maximal if the stimulated afferents and the stimulated hand muscle targeted with TMS are close to each other.

When considering the recency effect, heterotopic but not homotopic sensorimotor stimulation in the previous trial reduced the unconditioned MEP amplitude elicited in the subsequent trial. These findings indicate a reversed somatotopy of the recency effect compared to the known somatotopy of SAI shown in Fig. 1. A center-surround expression of inhibition, referring to the suppression of excitability in adjacent areas [39], could account for this reversal of the somatotopic gradient. Hence, heterotopic afferent stimulation of the little finger may trigger longer lasting surround inhibition of the central representation of the FDI muscle and therefore suppress the subsequent unconditioned response.

In addition, we cannot reject the possibility that the observed suppression of MEP responses following a heterotopically conditioned trial is driven by a shift in spatial attention. However, we consider this possibility to be unlikely because a potential shift in spatial attention also would decrease homotopic SAI, which we did not observe [40]. To the best of our knowledge only one previous study examined the effect of spatial attention towards a homotopic versus heterotopic skin area overlying the recorded hand muscles [41]. In that study, there was no effect of directing the attention to the skin overlying the abductor digiti minimi muscle when recording MEPs from the FDI muscle (i.e., attention directed to heterotopic muscle). In contrast, the MEP amplitude of FDI muscle was increased when attending to the skin above the FDI muscle. These results suggest that the reduction in MEP amplitude in the FDI muscle in trials following heterotopic stimulation of the little finger was not caused by a shift in attention towards the little finger. On the other hand, a homotopic facilitatory effect of attention on MEP amplitude may explain why conditioned trials with afferent stimulation of the index finger did not produce a consistent suppressive recency effect on MEPs recorded from FDI in the next trial.

### Limitations

As pointed out above, caution should be taken when concluding on the mechanisms underlying the observed effect of stimulation context. First, the experimental setup does not allow us to conclude whether it is the combination of sensory input and motor output or merely the sensory input that generates the recency effect. However, we dissociated a general recency effect of the previous MEP amplitude from a recency effect coupled to afferent conditioning during the randomized stimulation pattern, suggesting that a suppressive effect of peripheral nerve stimulation caused the recency effect. Second, in the randomized stimulation pattern the evaluation of the recency effect was carried out as a follow-up analysis. Consequently, we did not control the number of shifts between stimulation conditions, and the number of shifts were therefore not fully balanced within or between subjects. Lastly, the study consistently showed that the relative magnitude of SAI was decreased when conditions were randomly intermixed. However, it remains to be clarified if this effect is mediated at a cortical or subcortical level, or a combination of both.

## Abbreviations

TMS: transcranial magnetic stimulation
SAI: short-latency afferent inhibition
MEP: motor evoked potential
FDI: first dorsal interosseus muscle
MRI: magnetic resonance imaging
M1: primary motor cortex
D2: index finger
D5: little finger
SSEP: somatosensory evoked potential
EMG: electromyography
LMM: linear mixed-effects model

## Conclusion

We provide evidence that the pattern in which conditioned and unconditioned stimuli are intermingled affects the unconditioned MEP amplitude and thereby the relative magnitude of SAI. Further work is needed to elucidate the neural mechanisms and consequences of the dynamic tuning of corticospinal excitability by the recent history of sensorimotor stimulation.

## Acknowledgements

We would like to thank Mikkel Malling Beck and Kristoffer Hougaard Madsen for help with the statistical analyses.

Lasse Christiansen holds a postdoc grant from the Lundbeck Foundation (Grant nr. R322-2019-2406). Hartwig R. Siebner holds a 5-year clinical professorship in precision medicine at the Institute of Clinical Medicine, Faculty of Health and Medical Sciences, University of Copenhagen, sponsored by the Lundbeck Foundation (Grant nr. R186-2015-2138) and is supported by a collaborative research grant from the Lundbeck Foundation (Grant nr. R336-2020-1035). Marie T. Bonnesen received a research-year scholarship for pre-graduate students sponsored by the Lundbeck Foundation and the Danish Society for Neuroscience.

## Appendix: Figures

