## Supplementary Figures for "Corticospinal excitability is influenced by the recent history of electrical digital stimulation: implications for the relative magnitude of short-latency afferent inhibition"

### Bonnesen et al., Supplementary Material

#### Supplementary Figure 1

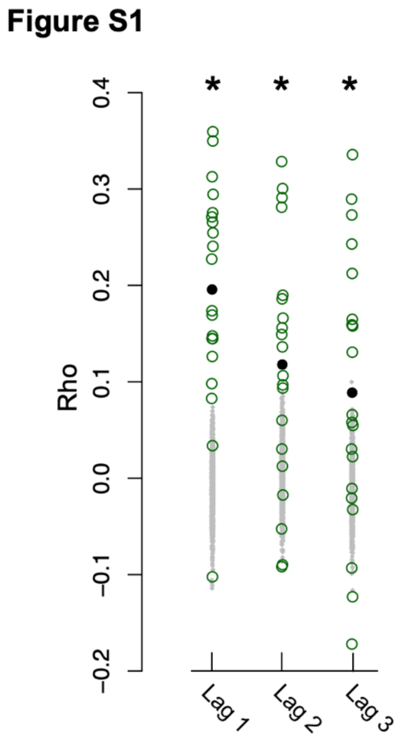


**Figure S1: Correlation coefficient between MEP response amplitude in each trial and response magnitude in the preceeding trials for data acquired with *Random* pattern.** Data were not divided into different conditions for this figure. Green circles depict individual Rho-values. Black dots reflect group-average. Gray dots reflect correlations computed for phase scrambled response vectors for each lag. * Denotes significant difference between correlation coefficients computed from trials within *Random* pattern and from phase-scrambled data.

#### Supplementary Figure S2-S10

#### Figure S2

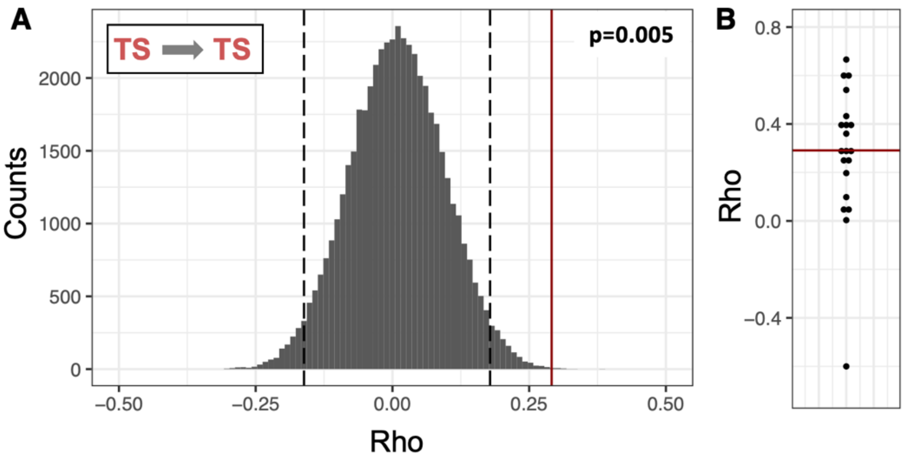


#### Figure S3

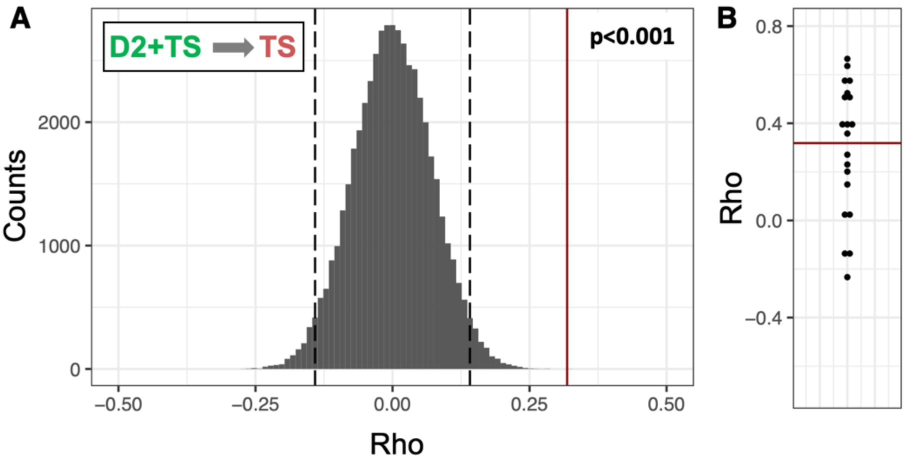


#### Figure S4

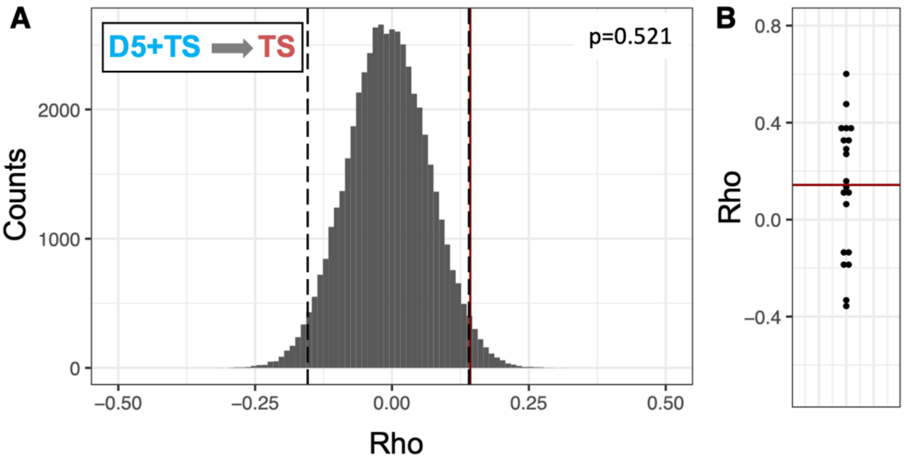

#### Figure S5

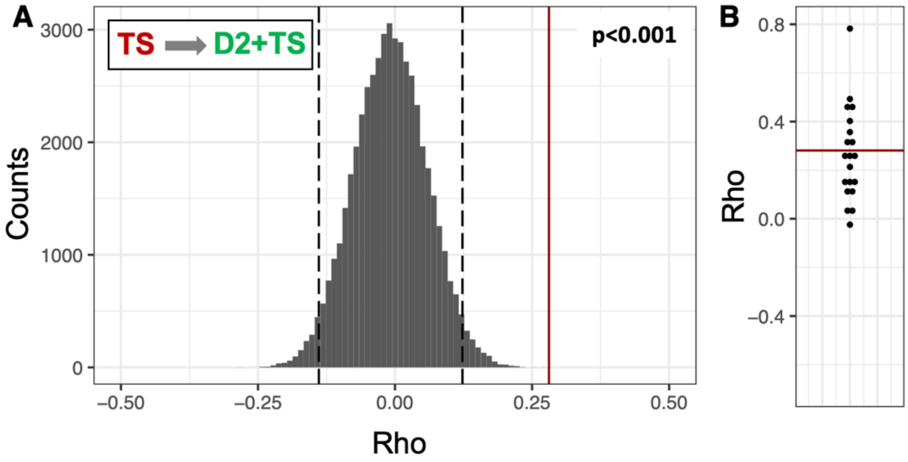


#### Figure S6

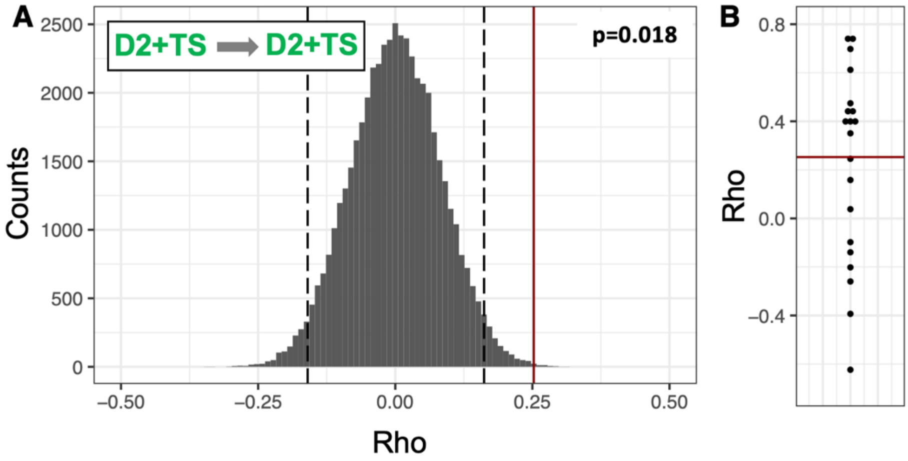


#### Figure S7

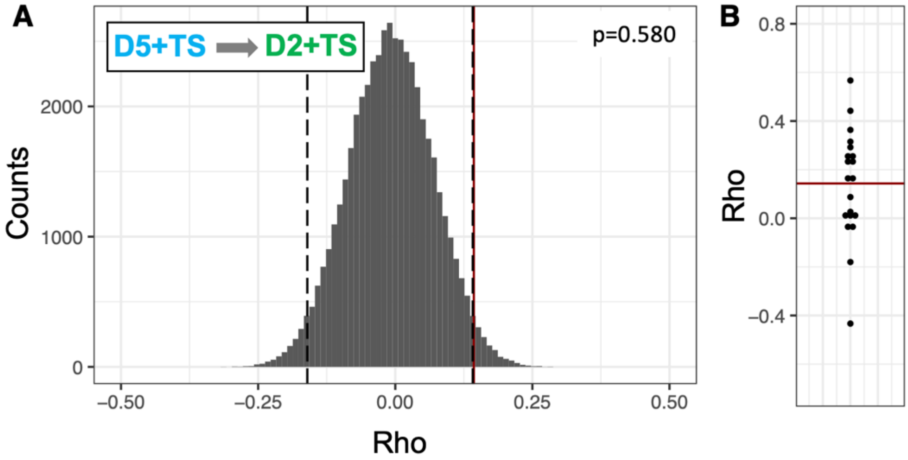

**Figure S8**

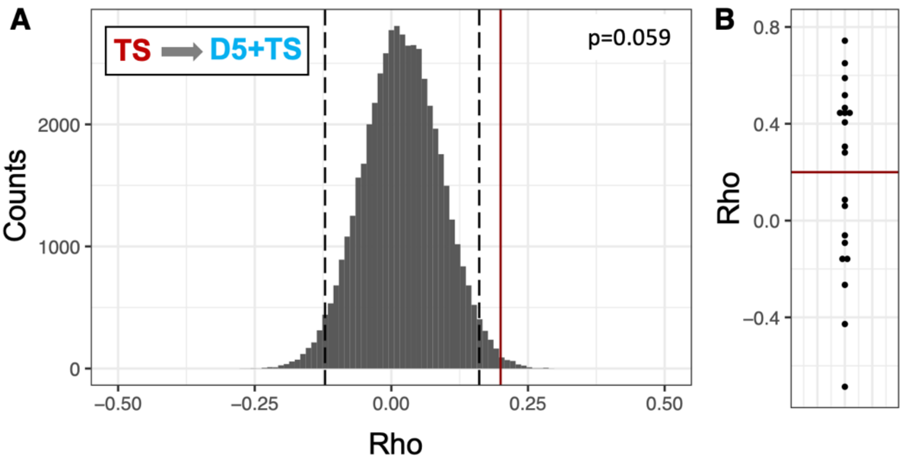


**Figure S9**

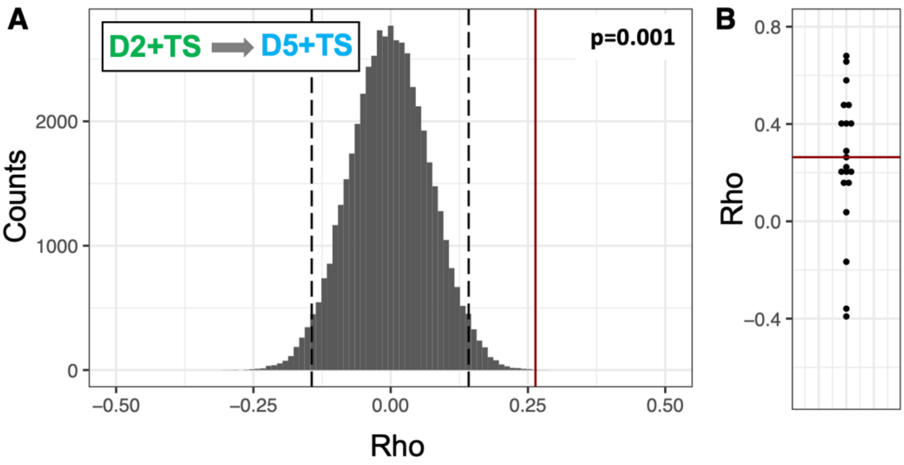


**Figure S10**

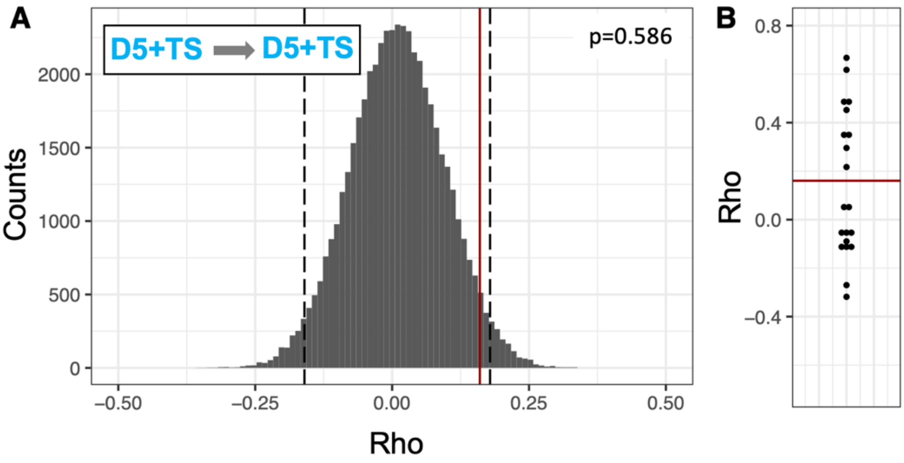

**Figure S2-S10: Correlation coefficient between MEP response amplitude in a given trial and response magnitude in the preceeding trial within and between conditions for data collected with *Random* pattern**.

**A)** The histograms reflect phase scrambled noise distribution. Columns (width of 0.1) depict the distribution of Rho-values from the 50000 correlations on phase-scrambled data. Vertical dashed lines depict 2.5% and 97.5% quantiles. The vertical, red line depicts group average.

**B)** Individual Rho values. Each dot reflect data from a single subject. The solid red line depict group-mean average. Averaging was done after Fisher’s Z-transformation and the average was subsequently inverse Fisher’s Z-transformed.
